# The Sequential Multispecies Coalescent

**DOI:** 10.1101/2025.01.31.635964

**Authors:** Analisa Milkey, Ming-Hui Chen, Yu-Bo Wang, Aolan Li, Paul O. Lewis

## Abstract

The multispecies coalescent (MSC) model applies coalescent theory to gene evolution within and among reproductively isolated populations (“species”) to estimate a species tree in the face of gene tree conflict resulting from deep coalescence. Sequential Monte Carlo (SMC) uses particle filtering to sample a posterior distribution, providing a fully-Bayesian and easily parallelized alternative to traditional MSC tree inference approaches. The method we propose samples first from the joint posterior distribution of gene and species trees, then samples species trees conditional on gene trees sampled previously, employing SMC for both rounds. Analyses of simulated and empirical datasets yield results comparable to state-of-the-art Bayesian MCMC approaches. Sampling the multispecies coalescent using SMC retains the advantages of fully Bayesian methods and is parallelizable in ways that Bayesian MCMC methods are not but also adds unique challenges. We demonstrate the performance of SMC compared to other commonly-used species tree methods using two empirical datasets and 400 simulated datasets.

---

Phylogenetic trees are essential tools in evolutionary biology, and many conclusions rest on their accuracy. Gene histories can differ from species histories due to deep coalescence, paralogy, horizontal gene transfer, and gene tree estimation error, which means many genes are necessary to accurately estimate the history of a set of species (Maddison, 1997) and, ideally, modeling should account for all sources of conflict. The multispecies coalescent model (MSC) (Rannala and Yang, 2003) accounts for deep coalescence (Pamilo and Nei, 1988), a major source of gene tree conflict, applying coalescent theory (Kingman, 1982) to explain the evolution of genes within populations constrained by the history of the reproductively-isolated units (species) to which they belong. In the most common version of the MSC, “species” are defined as completely reproductively isolated populations, so the “species tree” inferred is better termed a population structure tree (Sukumaran and Knowles, 2017); nevertheless, in this paper we will use the terms species and species tree to describe the reproductively-isolated units inferred by the MSC.

Under the basic Wright-Fisher population model (Fisher, 1930; Wright, 1931), generations are non-overlapping, mating is random (including a random amount of selfing), population size (*N* diploid individuals; 2*N* genes at any given locus) is constant through time, and different loci are unlinked and thus evolve independently. When multiple copies of a given gene are passed along to the next generation, the copying is termed a *coalescence event* when viewed looking backwards in time.

Deep coalescence occurs when two gene lineages sampled from a single species fail to coalesce before that species’ origin. Deep coalescence can result in the topology of the gene tree differing from the topology of other gene trees (gene tree conflict) as well as the species tree (Fig.1).

**Figure 1:**
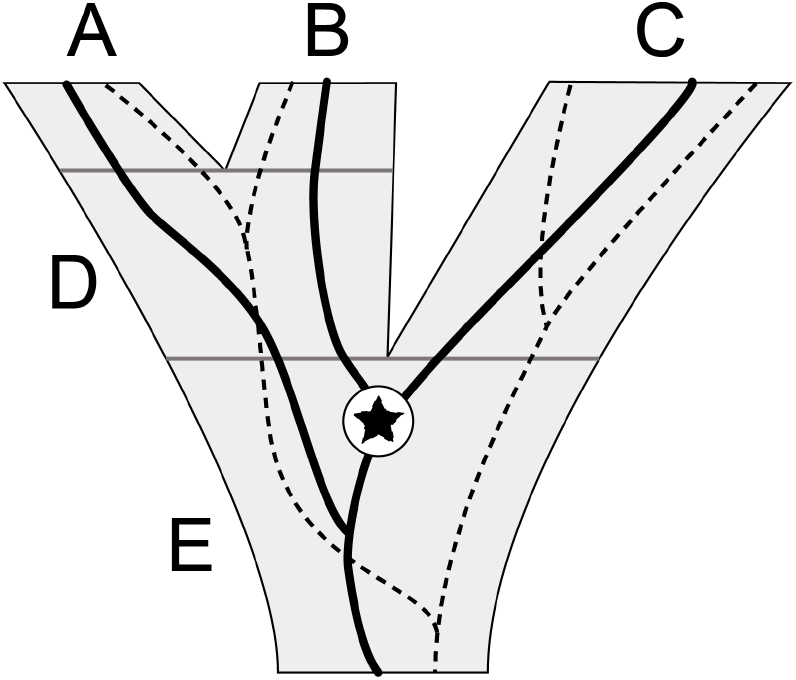
Gene trees for locus 1 (solid black) and locus 2 (dashed lines) embedded within a species tree (shaded). The species tree comprises 3 extant species (A, B, and C) and two ancestral species (D and E). Species boundaries are indicated by horizontal gray bars. The 2 lineages entering at the top of species D in locus 1 failed to coalesce until species E, illustrating deep coalescence (star), which results in the gene tree topology for locus 1 (A,(B,C)) conflicting with the topology ((A,B),C) of both the species tree and the gene tree for locus 2.

While fully Bayesian implementations of the MSC have considerable advantages (for example, estimation of gene trees and effective population sizes), they are computationally intensive and struggle to achieve convergence to the stationary posterior distribution on large data sets. StarBEAST3 is a fully Bayesian program that uses Markov Chain Monte Carlo (MCMC) to sample a multispecies coalescent posterior distribution (Heled and Drummond, 2010; Ogilvie et al., 2022; Douglas et al., 2014). Widely used alternatives include ASTRAL (Mirarab and Warnow, 2015), a summary method that takes estimated gene tree topologies as input and estimates the species tree topology from the frequency of different quartets of taxa. SVDQuartets, like ASTRAL, estimates only the species tree topology but has the advantage of taking into account uncertainty in gene trees while still being very computationally efficient (Chifman and Kubatko, 2014). Non-parametric bootstrapping of the original data can be performed with either ASTRAL or SVDQuartets to assess clade confidence. Recently, progress has been made on estimating speciation times in addition to topology using quartet methods (Peng, Swofford, and Kubatko, 2022), and local posteriors based on the MSC model provide support values for each edge of an ASTRAL species tree (Sayyari and Mirarab, 2016).

MCMC based on the Metropolis-Hastings algorithm is widely used in Bayesian phylogenetics and begins with a complete state (i.e., fully resolved phylogenetic tree with specified edge lengths) and proceeds by proposing local perturbations of the current tree as it runs. MCMC algorithms are challenging to parallelize because each iteration in the algorithm conditions on the previous iteration. Typically, parallelized MCMC approaches to phylogenetics involve splitting up likelihood calculations across processors, placing independent runs on different processors, or placing differently heated chains on different processors in the case of Metropolis-coupled MCMC. StarBEAST3 additionally places updates of gene trees from different loci on different processors.

Sequential Monte Carlo (SMC) or, more specifically, POSET SMC (Bouchard-Côté, 2012, 2014), is a Bayesian alternative to MCMC that uses particle filtering to sample a posterior distribution. In contrast to MCMC, SMC builds up a tree (whose state is stored in a “particle”) sequentially, resampling particles at each step based on particle weights that measure improvement in likelihood relative to the previous step (Fig. 2). SMC is naturally parallelizable in several ways that are not possible using the Metropolis-Hastings algorithm.

**Figure 2:**
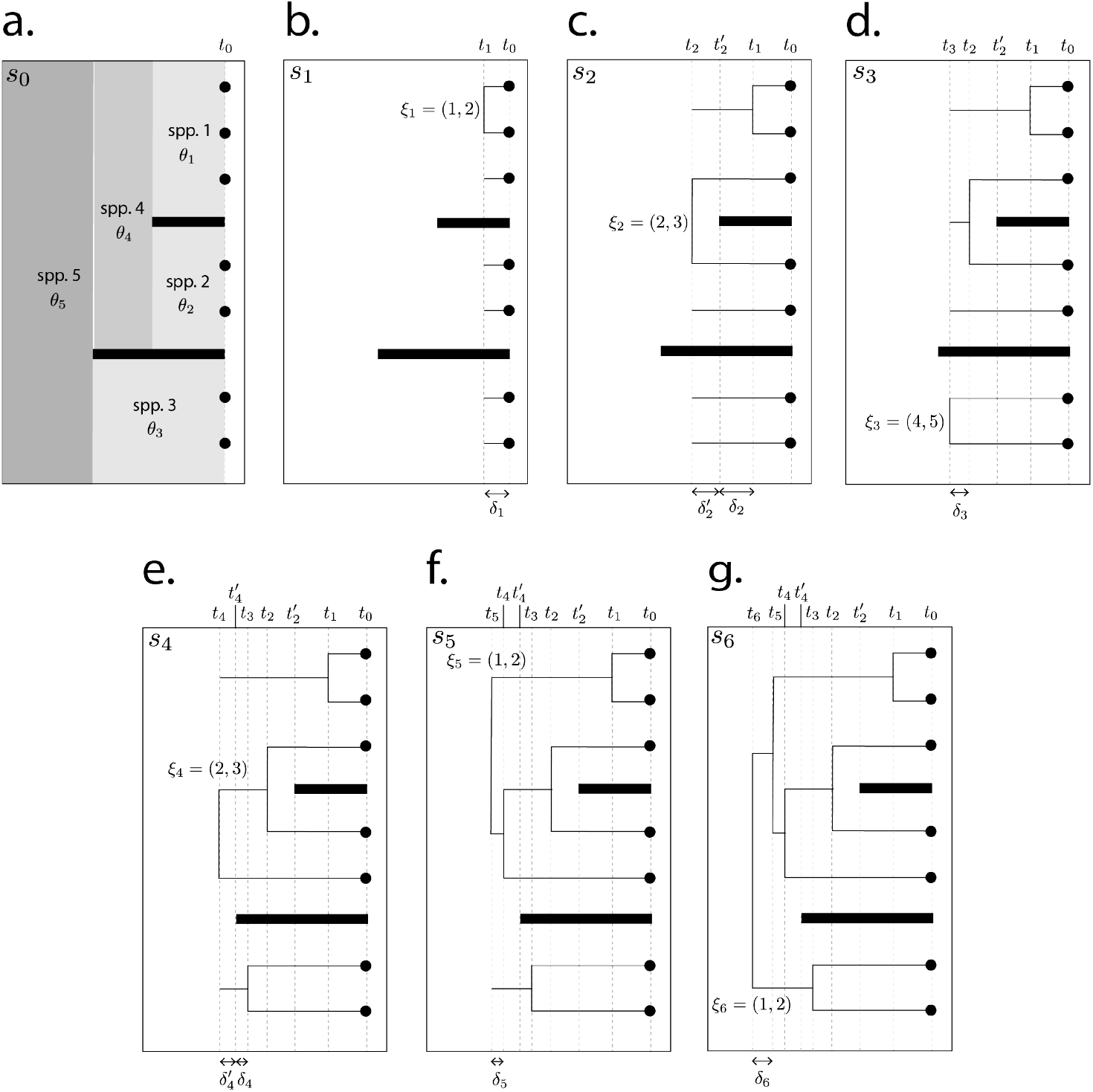
Species tree and growth of gene forest for one locus in one particle. Thick black lines represent species tree barriers to gene flow. Notation simplified by omitting locus and particle subscripts.

The filtering steps involved in SMC approaches are analogous to natural selection, and selective sweeps often occur, resulting in one particle’s “genome” replacing that of all (or nearly all) other particles, resulting in a condition known as particle degeneracy. Such sweeps lead to low effective sample size (ESS), analogous to the low ESS caused by autocorrelation in MCMC analyses. Thus, while SMC and MCMC both have potential failings, they are sufficiently different approaches that it is worthwhile exploring how well SMC can compete with MCMC-based MSC.

The SMC approach to the MSC model that we describe here differs from existing non-MCMC methods (ASTRAL, SVDQuartets) in its ability to deliver a sample from the posterior distribution of species trees (rather than a single point estimate) while sharing with these methods the ability to scale to datasets with many loci. Like Metropolis-Hastings MCMC methods, SMC is an approximation whose accuracy depends on the number of particles and the nature of the proposal distributions used. While Bouchard-Côté (2012) showed that, given unlimited computational resources, Metropolis-Hastings approaches can achieve higher accuracy than SMC, SMC can deliver reasonable results on a more practical time scale.

## Materials and Methods

Our approach employs SMC hierarchically. At the first level, SMC is used to obtain a sample from the joint posterior distribution of the species tree and gene trees from all loci. Species tree marginal distributions resulting from joint estimation typically suffer from particle degeneracy, and thus we employ a second level SMC to sample from the species tree posterior distribution conditional on the gene trees sampled during the lower-level SMC.

We first describe sampling from the joint posterior distribution (first level SMC) in the section entitled *SMC for Joint Gene and Species Tree Sampling*, then discuss the sampling from the conditional posterior (second level SMC) in the section *SMC for Species Trees Given Gene Trees*.

### SMC for Joint Gene and Species Tree Sampling

The joint posterior distribution for the simplest multispecies coalescent model (i.e., constant population size and speciation rate) can be written:

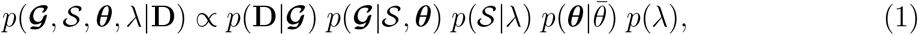

where:

**D** is a vector of aligned sequence data sets {**D**_*l*_ : *l* = 1, …, *L*} for *L* loci. Data set **D**_*l*_ comprises sequences of length *n*_*l*_ sites from each of *T* sampled genes from locus *l*;

***θ*** is a vector of mutation-scaled population size parameters (Watterson, 1975) {*θ*_*b*_ : *b* = 1, …, 2*M* − 1}, where *M* is the number of extant species (*M* ≤ *T*). Each edge *b* of species tree 𝒮 is associated with one element *θ*_*b*_ = 4*N*_*b*_*µ*_*b*_ of this vector, where *N*_*b*_ and *µ*_*b*_ are the effective (diploid) population size and mutation rate, respectively, for edge *b*;

***𝒢*** is a vector of gene trees ***𝒢***= {***𝒢***_*l*_ : *l* = 1, …, *L*}, where each component gene tree ***𝒢***_*l*_ comprises *T* − 1 increments and joins: ***𝒢***_*l*_ = {(*δ*_*t*_, *ξ*_*t*_) : *t* = 1, …, *T* − 1};

𝒮 is the species tree, comprising *M* − 1 increments and joins: 𝒮 = {(Δ_*m*_, Ξ_*m*_) : *m* = 1, …, *M* − 1};

*p*(**D**|***𝒢***) is the product of *Felsenstein likelihoods* (Felsenstein, 1981) computed using a substitution model (e.g., JC69; Jukes and Cantor, 1969) on each gene tree in ***𝒢***;

*p*(***𝒢***|𝒮, ***θ***) is the *coalescent likelihood* (Rannala and Yang, 2003), which is the probability density of the vector of gene trees ***𝒢*** conditioned on species tree 𝒮 and ***θ***;

*p*(𝒮|*λ*) is the *species tree prior*, assumed in this paper to be a pure-birth Yule model (Yule, 1925), where *λ* is the birth rate;

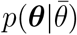 is the prior for the value of *θ* associated with each individual species, conditional on the assumed mean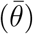; and *p*(*λ*) is the prior on the speciation rate *λ*, i.e., the single parameter of the Yule model.

### Proposing new states

The term *forest* is used for partial states comprising sets of disjoint, ultrametric subtrees, with each subtree having its own root vertex. The disjoint union of the leaves of the component subtrees yields the set of all sampled genes (for gene forests) or extant species (if the species forest). The term *tree* is reserved for complete states (i.e., a forest composed of just one subtree).

Each particle *k* (*k* = 1, …, *K*) begins with a vector of trivial gene forests ***𝒢***_*k*_ = {***𝒢***_*kl*_ : *l* = 1, …, *L*}. A trivial forest (state *s*_0_) comprises only leaf nodes and has height 0.0. A value for 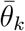 is drawn from its prior distribution. For each particle *k*, a complete species forest S_*k*_ is drawn from its Yule prior. For each edge *b* in S_*k*_, a mutation-scaled population size parameter value *θ*_*kb*_ is drawn from an InverseGamma (2,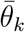) prior (Fig. 2a).

The remainder of this section describes proposing state *i* given state *i* − 1 in the gene forest for locus *l* in particle *k*; however, the subscript *k* is omitted to simplify notation. Each step *i* results in state *i* having one more coalescence event than state *i* − 1. Proposing one coalescence event for all *L* loci in a single particle thus involves *L* steps. Loci are visited in randomized order in each round of *L* steps.

Let *t* = *t*_*i*−1_ be the time at the start of step *i* (*i >* 0). A time increment *δ*_*li*_ is drawn from the coalescent prior distribution for locus *l*, which has rate *r*_*li*_,

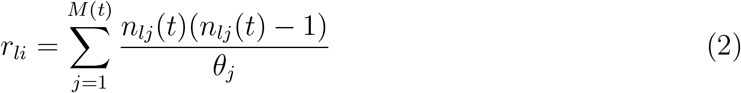

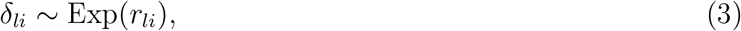

where *n*_*lj*_(*t*) is the number of uncoalesced gene tree lineages in existence at time *t* in species *j* and at locus *l*, and *M* (*t*) is the number of species at time *t*.

Let *τ*_1_ *<* … *< τ*_*M*−1_ be the heights of nodes representing speciation events in the species tree, and let 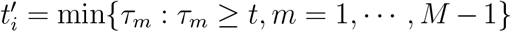. If 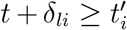 (e.g., Fig. 2c,e), then all lineages in gene forest *l* are advanced to time 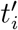 and another increment 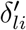 is drawn from an Exponential distribution whose rate now reflects the gene forest lineages merged as a result of the speciation event at time 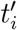. If 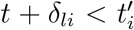 (e.g., Fig. 2b,d,f,g), all lineages in gene forest *l* are advanced by an amount *δ*_*li*_ and the species in which the next coalescent event occurs is determined using a draw from the multinomial probability distribution having parameters

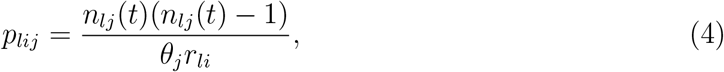

where *j* = 1, …, *M* (*t*) and *r*_*li*_ is the normalizing constant.

If species *j* is chosen, then two lineages *ξ*_*li*_ from species *j* in gene forest *l* are selected randomly from a Discrete Uniform distribution and joined. The new gene forest ancestral node is assigned to the same species as its two descendant lineages. The construction of state *i* is now complete.

Each step of this joint estimation SMC algorithm results in a new state. The number of steps *S* (and thus states) is thus the total number of coalescent events over all gene trees:

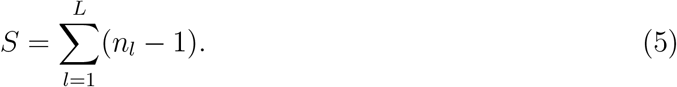

### Particle weights

Particle weights are calculated as the ratio of the product of the gene forest likelihoods after a coalescent event to the product of the gene forest likelihoods before a coalescent event (all prior terms in the numerator cancel with proposal terms in the denominator, leaving only the likelihood ratio).

The weight *w*_*i*_ for any given particle at state *s*_*i*_ (*i >* 0) is

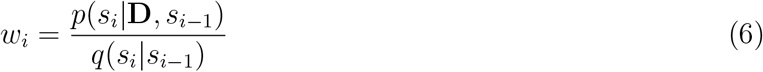

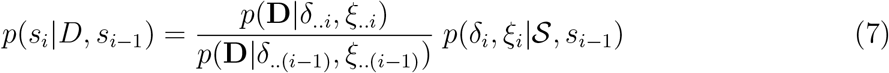

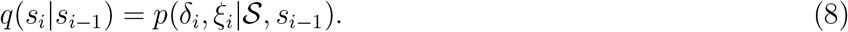

Note that we have suppressed the particle index *k* from the notation for readability, and use the shorthand notation *x*_..*i*_ = *x*_1_ … *x*_*i*_ for products of similar terms. The weight can be viewed as an importance weight in the context of importance sampling, where the conditional prior *q*(*s*_*i*_|*s*_*i*−1_) represents the importance density (Bouchard-Côté, 2014).

An exception is *w*_1_, which must take into account the prior probability of the species tree, which was simulated from the Yule prior when particles were first initialized:

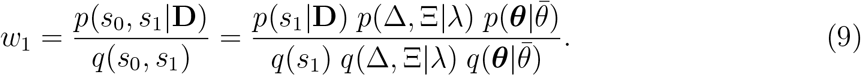

Because of the cancellation of prior terms with proposal terms, the weight *w*_*i*_ simplifies to

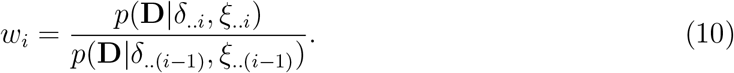

### UPGMA gene tree completion

The particle weight used in practice differs from (10) in one significant way. The weight defined in (10) sometimes leads to poor choices in the particle filtering stage (see below) because it lacks foresight. That is, a coalescence event proposed at step *i* may lead to a large improvement in likelihood at step *i*, but, in the final gene tree, results in a lower likelihood than if a different join had been made at step *i*. Bouchard-Côté (2012) described an alternative proposal where trees in a partial state are (temporarily) completed using a fast, deterministic approach such as neighbor-joining (NJ). After the weight is calculated using this temporary complete state, the portion of the tree completed using NJ is removed.

We implement this approach using a proposal where gene forests are completed using a UPGMA algorithm (Sokal and Michener, 1958) with Jukes-Cantor distances (Jukes and Cantor, 1969). We use UPGMA rather than NJ to maintain ultrametricity. The likelihoods in both numerator and denominator of the particle weight are thus based on complete gene trees in which state *s*_*i*_ is embedded. This allows SMC to “look ahead” and leads to better-informed choices at each step. The weight including this modification is

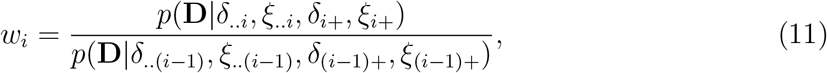

where *δ*_*i*+_ and *ξ*_*i*+_ are the increments and joins added to state *i* using the UPGMA algorithm and *δ*_(*i*−1)+_ and *ξ*_(*i*−1)+_ are the increments and joins added to state *i* − 1 using the UPGMA algorithm.

### Filtering

Filtering of particles takes place after a coalescent event has been proposed in each particle for a given locus. Filtering is performed using multinomial sampling with normalized weights as bin probabilities. If a particle is selected, its species tree 𝒮 (including ***θ***) as well as its vector of gene forests ***𝒢*** is copied to the new particle generation.

A selective sweep may result in one particle (with its associated species tree) replacing all other particles. At this point, the part of the species tree beyond the deepest coalescent event in any locus has not been influenced by the data, yet all future proposals will be constrained by this species tree. Thus, after each locus has undergone filtering, the species tree in each particle is trimmed back to the deepest coalescent event across all loci in that particle and rebuilt from that point on, with new values of *θ* drawn for each new species population. This reintroduces variation in species trees across particles.

Why is filtering performed after gene forests in a single locus have been updated rather than after all loci have been considered? If loci differ in sequence length, a locus with a longer sequence will often achieve a larger difference in likelihood compared to a locus with a shorter sequence length. Filtering loci independently ensures that loci with longer sequences do not exert undo dominance in determining particle populations, especially at early stages.

### SMC for Species Trees Given Gene Trees

The first-level SMC described in the previous section often results in a marginal posterior species tree sample that is dominated by very few distinct species trees. This is because the Felsenstein likelihood is only indirectly influenced by the species tree hyperparameter through its effect on gene tree edge lengths and joins. In contrast, the Felsenstein likelihood directly affects the filtering of gene forests at each stage. We thus resample the marginal species tree posterior in the second-level SMC, conditioning on the gene trees sampled in the first-level SMC.

Using SMC to estimate the distribution of species trees conditional on complete gene trees and 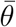 (mean mutation-scaled population size *θ*) is very similar to the method described in detail by Bouchard-Côté (2012) and Bouchard-Côté (2014), with the primary difference being that the likelihood function is the integrated coalescent likelihood (Jones, 2017) rather than the Felsenstein (1981) likelihood.

If complete gene trees are available, the conditional posterior distribution of species trees does not require calculation of the Felsenstein likelihood,

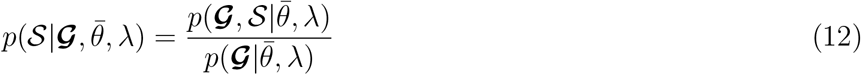

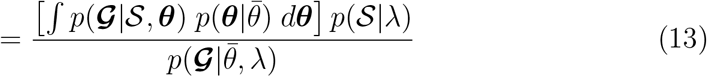

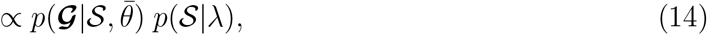

Where 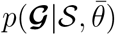 is the *integrated coalescent likelihood* (Jones, 2017). Because this distribution involves only the integrated coalescent likelihood and not the Felsenstein likelihood, sampling is much less computationally demanding.

### Increments and joins

The second-level SMC begins with *K*^*^ particles, each having *L* complete gene trees and a species forest in the trivial state (*s*_0_), consisting of only the *M* leaf vertices, each having height 0.0 (Fig. 3a). The vector of gene trees is sampled randomly from the first-level particle population and is shared by every particle in the second-level analysis.

**Figure 3:**
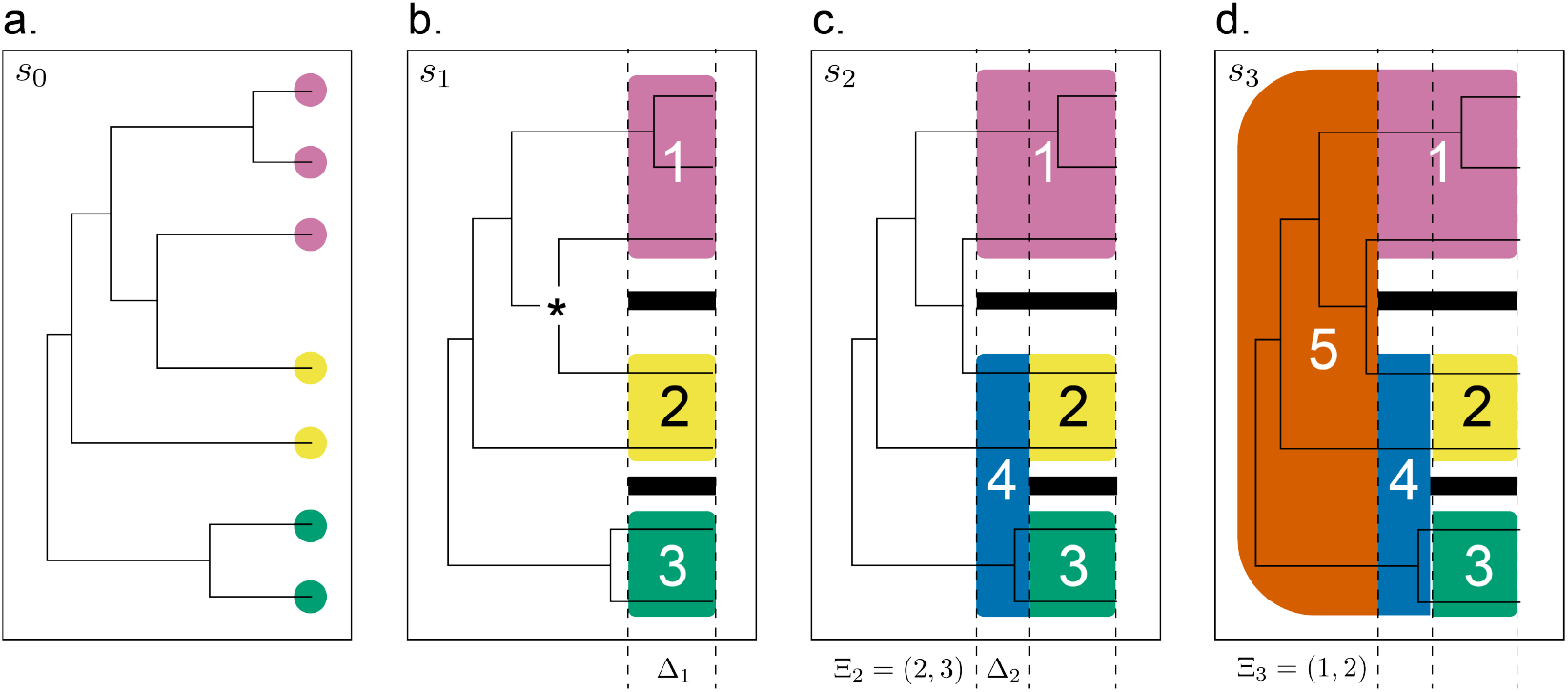
Growth of species forest (*M* = 3 species), constrained by a gene tree from one locus in a single particle. a. trivial state *s*_0_. b. partial state *s*_1_. c. partial state *s*_2_. d. complete state *s*_3_. Black bars represent reproductive isolation barriers separating distinct species, numbers indicate distinct species, and the asterisk (*) denotes the coalescence event that marks the maximum possible value of Δ_1_ (and also Δ_2_ given that Ξ_2_ joined species 2 and 3).

For example, a first-level analysis might use *K* = 1000 particles, of which 10% are sampled for use in the second-level analysis. If *K*^*^ = 200 particles are used for the second-level analysis, then a total of (0.1*K*)*K*^*^ = (0.1 ^*^ 1000) ^*^ 200 = 20000 species trees would constitute the second-level posterior sample, with each of 0.1*K* = 0.1 ^*^ 1000 = 100 gene tree vectors retained from the first-level SMC forming the basis for an independent second-level SMC involving *K*^*^ particles.

The state of every particle is advanced from the trivial (species forest) state through a series of partial states to a complete state via a series of proposals. A weight is computed for a proposed new state and particles are filtered by drawing *K*^*^ particles with replacement from a multinomial distribution in which the bin probabilities are the normalized particle weights.

There are *j* = *M* − *i* + 2 lineages before state *i* is proposed (*i* = 2, …, *M*). The transition from partial state *s*_*i*−1_ to partial state *s*_*i*_ involves first choosing a pair Ξ_*i*_ of existing lineages to join, with probability 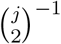, and then a height increment Δ_*i*_ from the Exponential (*jλ*) prior distribution:

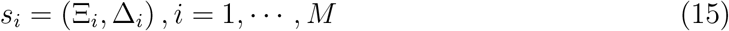

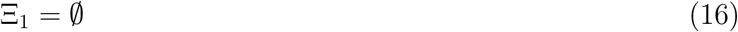

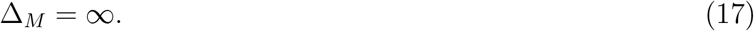

The first (*i* = 1) and last (*i* = *M*) steps represent exceptions: (1) two lineages are not joined in creating partial state *s*_1_ because there is no reason to assume that the most recent speciation event occurred exactly at time 0 and (2) the final increment is, necessarily, ∞ and thus is not a random variable; hence, Ξ_1_ is the empty set, and Δ_*M*_ = ∞. In all partial states *s*_*i*_ (*i < M*), we implicitly assume an *ephemeral* ancestral species, which extends from *τ*_*i*_ = _*j*≤*i*_ Δ_*j*_ to ∞. This is necessary because species forests are conditioned on complete gene *trees* (not partial-state gene *forests*).

To illustrate why Ξ_*i*_ is chosen before Δ_*i*_ in each step, consider state *s*_1_ in Figure 3b. The coalescent likelihood must account for 1 coalescent event in species 1 (top) and 0 coalescent events in both species 2 and 3 (middle and bottom) during the time interval (0, Δ_1_). It must also account for the remaining 5 coalescent events in the history of the 6 lineages that exist at time Δ_1_. These 6 lineages and their ancestors are all members of the ephemeral ancestral species. Joining two lineages after choosing the increment Δ_1_ would therefore have no effect on the coalescent likelihood because such a join would be associated with a time interval of zero length between the join and the start of the ephemeral species. The consequences of such a join would not be realized until the *next* step, at which point there is no longer any opportunity to make the particle pay for a poor join decision. Thus, joins always follow increments when constructing species forests in the second-level SMC.

In general, the number of loci is greater than 1, so the species forest constructed within each particle is conditioned on gene trees from more than one locus. The coalescent events within gene trees place constraints on the maximum value that any given species tree increment can attain. For example, the increment Δ_1_ in Figure 3b must be less than or equal to the time of the coalescence event indicated by the asterisk (*). This is the first coalescence event (over all loci) where lineages from two distinct species join. Extending Δ_*i*_ further back in time than this gene tree node would imply gene flow across species boundaries, which is not allowed under the multispecies coalescent model. Such constraints lead to sampling efficiencies even if the prior on species tree increments is vague.

### Particle weights

As for the first-level SMC, the weight *w*_*k*_ for particle *k* (*k* = 1, …, *K*) at state *s*_*i*_ can be viewed as an importance weight in the context of importance sampling:

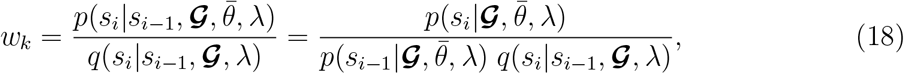

Where 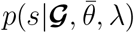 is the posterior probability density of state *s* and *q*(*s*_*i*_|*s*_*i*−1_, ***𝒢***, *λ*) is the importance density, which, in this case, equals the proposal density for Δ_*i*_ and Ξ_*i*_ given the previous state *s*_*i*−1_:

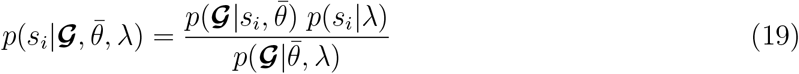

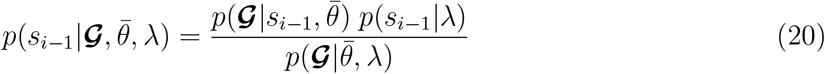

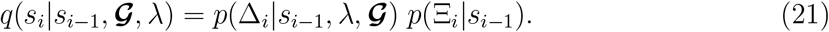

The multispecies coalescent likelihood can be computed piecewise by species tree edge. Figure 4 illustrates the calculation of the coalescent likelihood for locus *l* on edge *b* of the species tree. If edge *b* is one of the edges added to construct state *i*, then 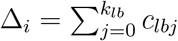.

**Figure 4:**
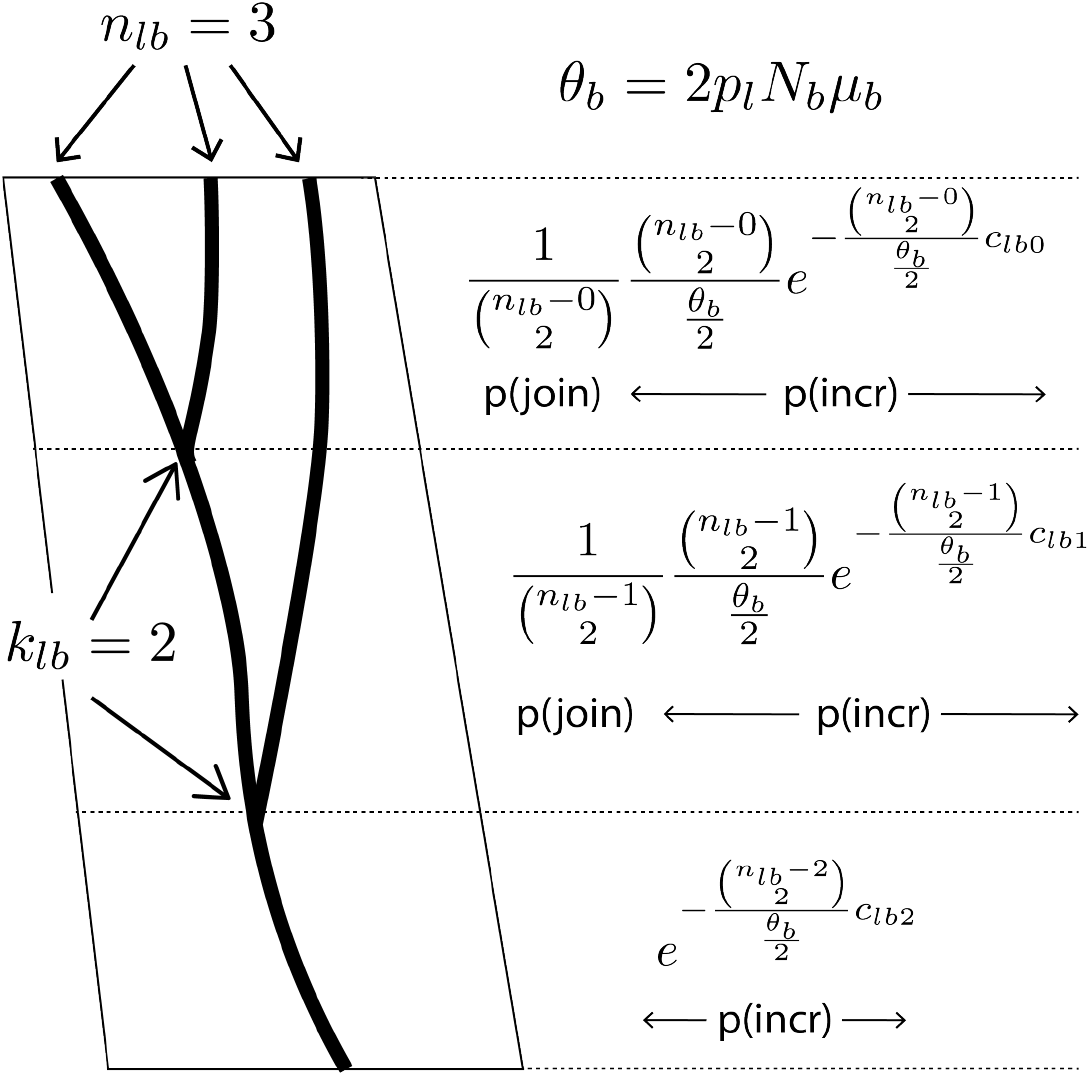
Computation of the multispecies coalescent likelihood for edge *b* of the species tree at locus *l*. The *k*_*lb*_ coalescent events partition the edge length into intervals of length *c*_*lb*0_, *c*_*lb*1_, and *c*_*lb*2_. *n*_*lb*_ and *n*_*lb*_ – *k*_*lb*_ are the number of lineages entering and leaving the edge, respectively.

Jones (2017) observed that the coalescent likelihood takes the form of an InverseGamma(*q*_*b*_ −1, *γ*_*b*_) distribution for each edge *b* of the species tree:

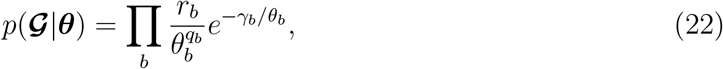

where

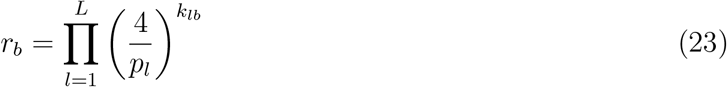

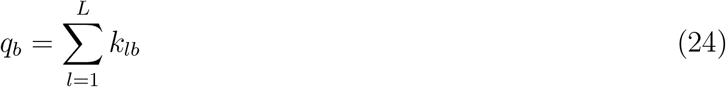

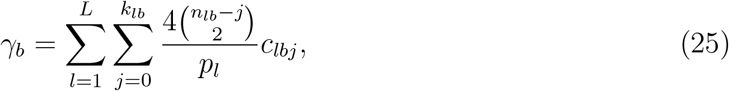

*b* indexes edges (i.e., species) in the species tree, *l* indexes loci, *i* indexes coalescent events within species *b* in gene tree ***𝒢***_*l*_, *p*_*l*_ is the ploidy level of locus *l* (e.g., *p*_*l*_ = 1 for a plastid or mitochondrial locus and *p*_*l*_ = 2 for nuclear loci in diploid organisms), *c*_*lbj*_ is the *j*th time interval in gene tree ***𝒢***_*l*_ for edge *b*, and *n*_*lb*_ is the number of lineages in gene tree ***𝒢***_*l*_ at the start of edge *b*. Note that our formulas for *r*_*b*_ and *γ*_*b*_ contain the factor 4 because the Watterson (1975) definition of *θ*_*b*_ that we use (*θ*_*b*_ = 4*N*_*b*_*µ*_*b*_) differs from Jones (2017), who used *θ*_*b*_ = *N*_*b*_*µ*_*b*_, where *N*_*b*_ is the effective population size and *µ*_*b*_ the mutation rate specific to edge *b*.

Assuming an InverseGamma(*α, β*) prior distribution for *θ*_*b*_ allows *θ*_*b*_ to be analytically integrated out of the coalescent likelihood (Jones, 2017):

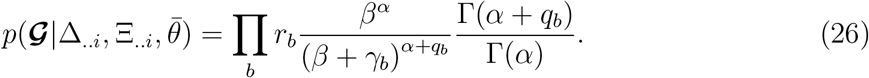

We assume *α* = 2, 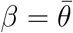, where the parameter 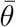 specifies the mean value of *θ* among species.

For the Yule pure-birth tree model, the proposal density for Δ_*i*_ is

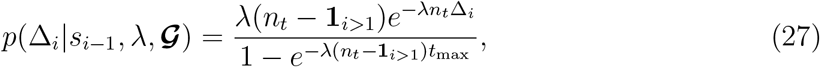

where *n*_*t*_ is the number of species tree lineages at time *t, t* = Σ_*j<i*_ Δ_*j*_ denotes the height of the species forest at state *s*_*i*−1_, *t*_max_ is the upper bound for Δ_*i*_ determined by ***𝒢***, and **1**_*i>*1_ is an indicator variable that is 1 if *i >* 1 and 0 if *i* = 1.

The join proposal probability is

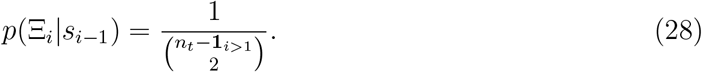

Cancellation of prior terms in the numerator with proposal terms in the denominator results in the following particle weight specification:

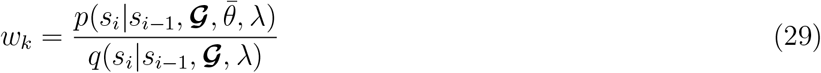

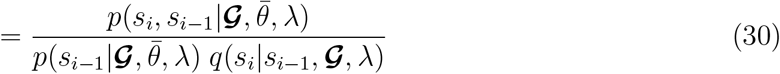

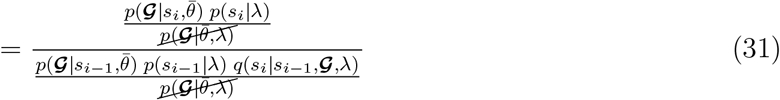

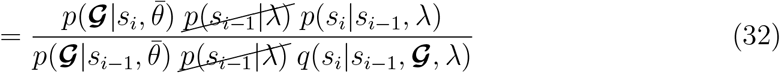

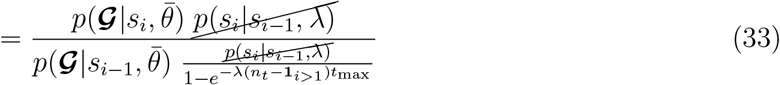

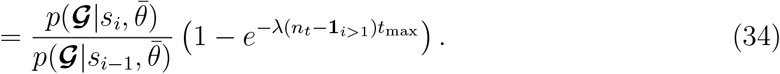

The first term is the ratio of the integrated coalescent likelihood for state *s*_*i*_ to the integrated coalescent likelihood for state *s*_*i*−1_. The second term is the normalizing constant of the truncated Exponential proposal distribution for increment Δ_*i*_.

### Particle filtering

After parameters Δ_*i*_ and Ξ_*i*_ are proposed and weights are determined for each particle, the weights are normalized and multinomial sampling is used to draw *K*^*^ new particles using the normalized weight 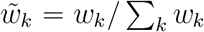 as the probability for bin *k*. After the filtering step, the particle population represents a sample from the posterior distribution of state *s*_*i*_ = {Δ_..*i*_, Ξ_..*i*_} conditional on 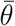 and gene trees ***𝒢***.

### Simulations

We performed simulations to compare the performance of the SMC approach to the Bayesian MCMC approach of StarBEAST3 with respect to accuracy of the species tree topology. At every point in a 20 by 20 grid, data were simulated for 10 loci, 5 species, and 2 sampled individuals per species. The 20 grid rows corresponded to evenly-spaced values of *T*, the species tree height, from 0.0 to 0.5. The 20 grid columns corresponded to evenly-spaced values of *θ/*2, from 0.0 to 0.3. *θ* was fixed across all species within a species tree. Smaller values of *T* and larger values of *θ/*2 yield greater expected deep coalescences, and thus those areas of the grid present more challenges to species tree methods.

Species trees were simulated using a pure-birth Yule (1925) model with speciation rate *λ*. Gene trees were simulated within the species tree from the prior. Sequences between 200 and 1000 nucleotides were simulated under the Jukes-Cantor substitution model with equal rates among sites and loci.

SMC analyses used *K* = 10000 particles for the first level and *K*^*^ = 500 particles for the second level. A randomly-chosen 2.5% of particles from the first-level analysis were used for the second-level analysis. A randomly-chosen 0.8% of second-level particles were saved, yielding a sample of size 1000 species trees. StarBEAST3 analyses used 15 million iterations, saving every 15000 for a total sample size of 1000 species trees. Burn-in was set to 1.5 million iterations.

Analyses using both SMC and StarBEAST3 assumed a fixed speciation rate determined from the tree length estimated by the QAGE method (Peng, Swofford, and Kubatko, 2022). Note that the simulations assumed *θ* was constant throughout the species tree, whereas both SMC and StarBEAST3 allowed *θ* to vary among species; however, both assumed constant *θ* within a species. The mean *θ* used by both SMC and StarBEAST3 analyses was also estimated by the QAGE method. The definition of *θ* differed between SMC and StarBEAST3. StarBEAST3 defines *θ* = *N*_*e*_*µ* whereas SMC assumes the Watterson (1975) mutation-scaled population size definition, *θ* = 4*N*_*e*_*µ*. To make analyses comparable, we thus set StarBEAST3 mean *θ* to 1/4 that of SMC.

The Robinson-Foulds (RF; Robinson and Foulds, 1981) distances between each sampled species tree and the true species tree used for simulation were averaged to provide a measure of species tree topology accuracy.

### Empirical Data

We also explored the performance of our method on two datasets, one containing gopher species and one containing snake species. The gopher dataset consists of a subset of data from Belfiore et al. (2008). The data contain 6 loci sampled from 27 individuals representing 10 species of pocket gophers. The 9 ingroup species belong to the genus *Thomomys*, and the outgroup belongs to the genus *Orthogeomys*. The snake dataset consists of a subset of data from Chifman and Kubatko (2014). The data contain 15 loci sampled from 52 individuals representing 7 species of snakes. The original data set contained 19 loci; we removed 4 loci in which one or more taxa had completely missing data. The 6 ingroup species belong to the genus *Sistrurus*, pygmy rattlesnakes, and the outgroup species belongs to the genus *Agkistrodon*. We chose these datasets because they are well-studied, making them good measures of program performance. They are also small enough that it is feasible for multiple programs, even computationally intensive programs, to run them in a reasonable amount of time.

### Gopher dataset

We estimated a species tree using StarBEAST3 (Douglas et al., 2014) and SMC. In both programs, we used a Jukes Cantor (Jukes and Cantor, 1969) substitution model and a fixed speciation rate determined from the tree length estimated by QAGE (Peng, Swofford, and Kubatko, 2022). Mean *θ* was also estimated by the QAGE method. We ran StarBEAST3 for 20 million generations, with 2 million generations of burnin, saving trees every 10000 generations for a total of 2000 species trees sampled. We allowed each locus to have a different relative rate of substitution with default prior LogNormal(1.0, 0.6). SMC analyses used *K* = 10000 particles for the first level and *K*^*^ = 1000 particles for the second level. A randomly-chosen 2.5% of particles from the first-level analysis were used for the second-level analysis. A randomly-chosen 0.8% of second-level particles were saved, yielding a sample of size 2000 species trees. We fixed relative rates for each locus according to estimates under a site-specific model implemented in PAUP*. We conducted 10 independent runs in both StarBEAST3 and SMC and combined the resulting species trees, yielding a sample 20,000 species trees from each program.

We also estimated species trees using SVDQuartets and ASTRAL IV (as implemented in ASTER (Zhang & Mirarab, 2022; Tabatabaee et al., 2023)). The SVDQuartets analysis comprised 1000 bootstrap replicates, and the tree was rooted at outgroup *O. heterodus*. For the ASTRAL analysis, we used maximum likelihood gene trees estimated from IQTREE (Nguyen et al., 2015), using a Jukes Cantor model for each locus. We rooted the tree at outgroup *O. heterodus*.

### Snake dataset

We estimated a species tree using StarBEAST3 (Douglas et al., 2014) and SMC. In both programs, we used a Jukes Cantor (Jukes and Cantor, 1969) substitution model and a fixed speciation rate determined from the tree length estimated by QAGE (Peng, Swofford, and Kubatko, 2022). Mean *θ* was also estimated by the QAGE method. We ran StarBEAST3 for 100 million generations, with 10 million generations of burn-in, saving trees every 50000 generations for a total of 2000 species trees sampled. We allowed each locus to have a different relative rate of substitution with default prior LogNormal(1.0, 0.6). SMC analyses used *K* = 15000 particles for the first level and *K*^*^ = 1000 particles for the second level. A randomly-chosen 1.67% of particles from the first-level analysis were used for the second-level analysis. A randomly-chosen 0.8% of second-level particles were saved, yielding a sample of size 2000 species trees. We fixed relative rates for each locus according to estimates under a site-specific model implemented in PAUP*. We conducted 10 independent runs in both StarBEAST3 and SMC and combined the resulting species trees, yielding a sample of 20,000 species trees from each program.

We also estimated species trees using SVDQuartets and ASTRAL IV (as implemented in ASTER (Zhang & Mirarab, 2022; Tabatabaee et al., 2023)). For the SVDQuartets analysis, we used 1000 bootstrap replicates and rooted the tree at outgroup *Agkistrodon spp*. For the ASTRAL analysis, we used maximum likelihood gene trees estimated from IQTREE (Nguyen et al., 2015), using a Jukes Cantor model for each locus. We rooted the tree at outgroup *Agkistrodon spp*.

## Results

### Simulations

Without accounting for parallelization, SMC took 1126 seconds on average, and StarBEAST3 took 1359 seconds on average. We chose settings that gave each program roughly the same computational budget to fairly assess accuracy.

As expected, smaller values of *T* combined with larger values of *θ/*2 produced difficult conditions, with more than 60% of the maximum possible number of deep coalescences being deep (Fig. 5a, lower right). Larger values of *T* combined with larger *θ/*2 yielded easier conditions (Fig. 5a, upper left).

**Figure 5:**
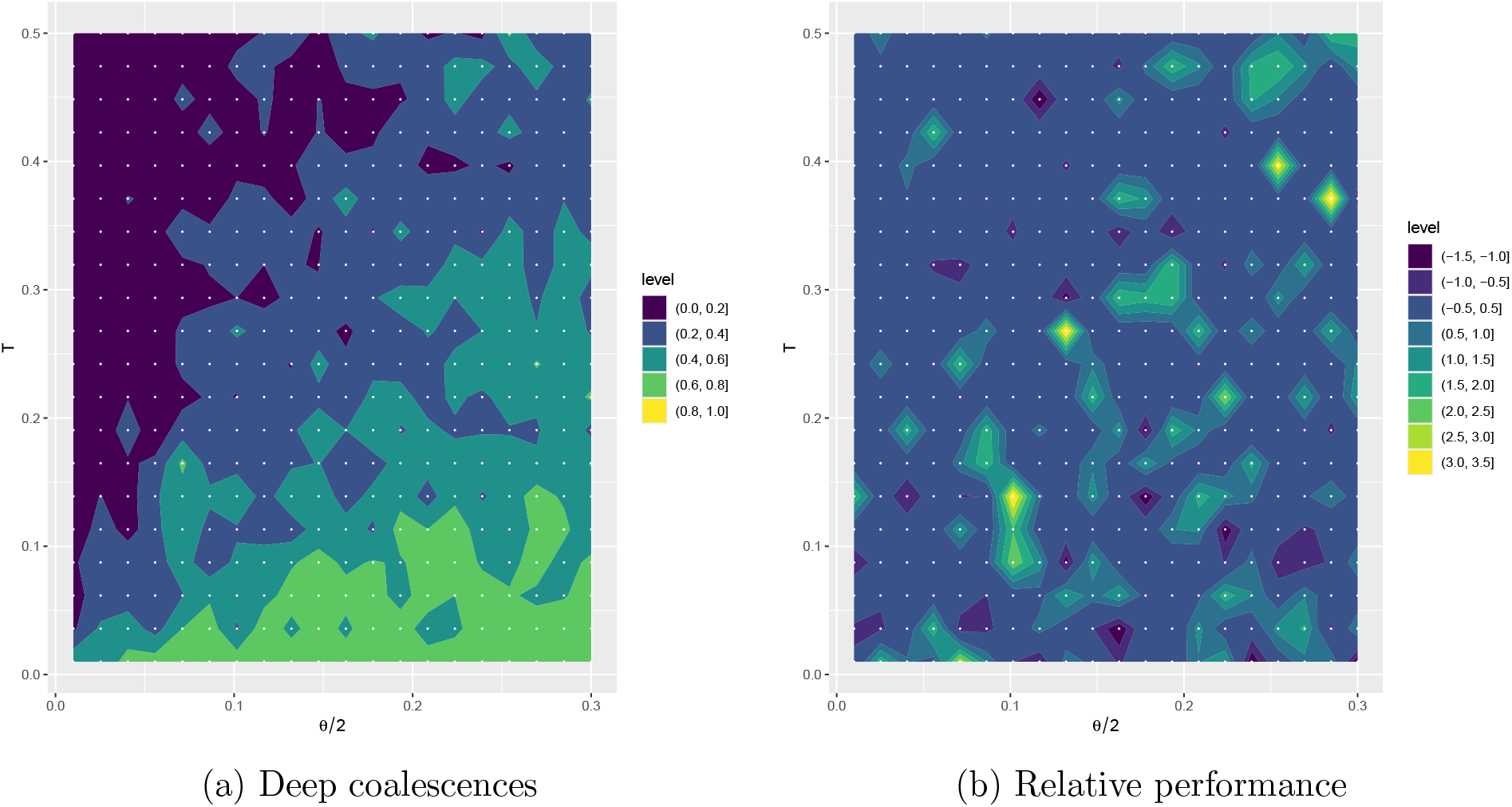
Plots show simulation results for 400 combinations of expected species tree height *T* and *θ/*2. a) Ratio of true number of deep coalescences to the maximum possible number of deep coalescences. (Higher values represent more difficult parameter combinations.) b) Mean SMC Robinson-Foulds (RF) distance to the true tree minus mean StarBEAST3 RF distance to the true tree. Positive values (yellow) indicate StarBEAST3 performed better for the simulation; negative values (purple) indicate SMC performed better. Points indicate (*θ/*2, *T*) combinations. Plot shows surface smoothed between sampled points.

SMC and StarBEAST3 performed similarly (average difference in RF distance to true species tree between -0.5 and 0.5) for the majority (270 out of 400) of combinations of *T* and *θ/*2 (Fig. 5b). There is no area of the grid where one method consistently outperforms the other. That said, StarBEAST3 performed slightly better than SMC on average, with an average RF distance of 0.399, compared to 0.638 for SMC. SVDQuartets produced an average RF distance of 1.005, and ASTRAL produced an average RF distance of 0.775.

### Empirical Data

*Gopher dataset*.*—* All four methods found support for the clade containing *T. townsendii, T. bottae*, and *T. umbrinus* (Fig. 6). SMC, StarBEAST3, and ASTRAL recovered *T. bottae* and *T. townsendii* as sister, while SVDQuartets recovered *T. townsendii* and *T. umbrinus* as sister. All methods also found support for the clade containing *T. idahoensis, T. monticola, T. mazama*, and *T. talpoides*. ASTRAL and SVDQuartets found support for *T. idahoensis* and *T. talpoides* as sister within this clade, with *T. mazama* as the most distantly related species within this clade. SMC and StarBEAST3 found *<*50% support for any sister-group relationship in this clade. Neither StarBEAST3 nor SMC found support for a sister-group relationship between the putative outgroup *O. heterodus* and the ingroup species, though trees are displayed rooted at this taxon. (Neither SVDQuartets nor ASTRAL infer the root.) Without accounting for parallelization, SMC took 2298 seconds on average, and StarBEAST3 took 2629 seconds on average.

**Figure 6:**
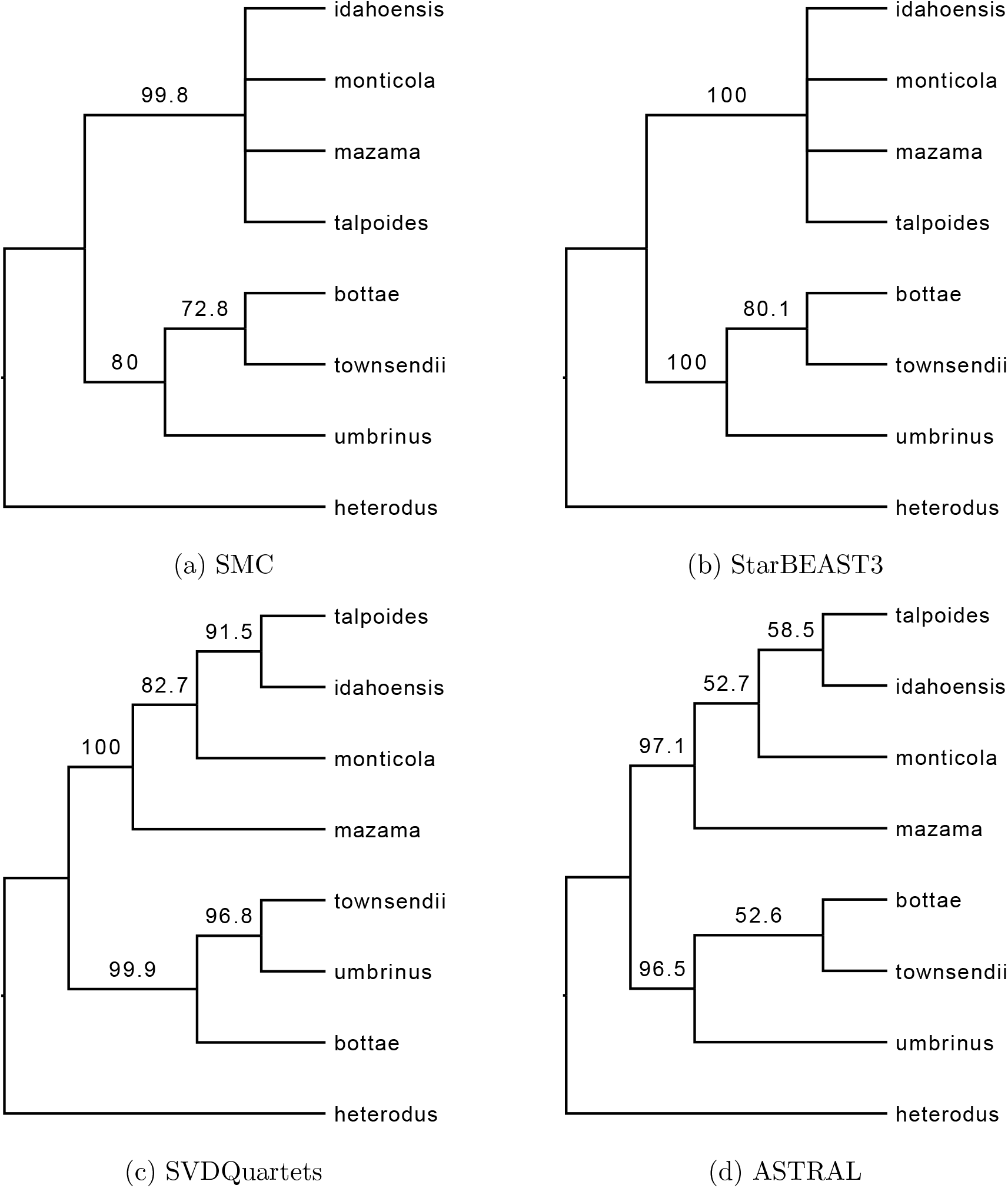
Gopher dataset. a) 50% majority rule consensus tree estimated by SMC. b) 50% majority rule consensus tree estimated by StarBEAST3. c) Bootstrapped consensus tree estimated by SVDQuartets. d) Tree estimated by ASTRAL with local posterior probabilities. All trees rooted at outgroup *O. heterodus*.

Maximum clade credibility (MCC) trees for SMC (Fig. 7a) and StarBEAST3 (Fig. 7b) show comparable branch lengths between programs. SMC found slightly longer branch lengths for the clade containing *T. townsendii, T. bottae*, and *T. umbrinus*, but branch lengths on the MCC trees for this clade are no more than 0.0026 units different. Posterior probabilities for the MCC trees (Fig. 7c,d) are also comparable. Although both programs found strongest support for different sister relationships within the *T. idahoensis, T. monticola, T. mazama*, and *T. talpoides* clade, posterior probabilities were no greater than 0.5 for any relationship in this clade. The MCC trees are *not* rooted at the putative outgroup *O. heterodus*. SMC found lower support than StarBEAST3 for this taxon in the clade containing *T. townsendii, T. bottae*, and *T. umbrinus*, placing more support for this taxon as sister to the putative ingroup.

**Figure 7:**
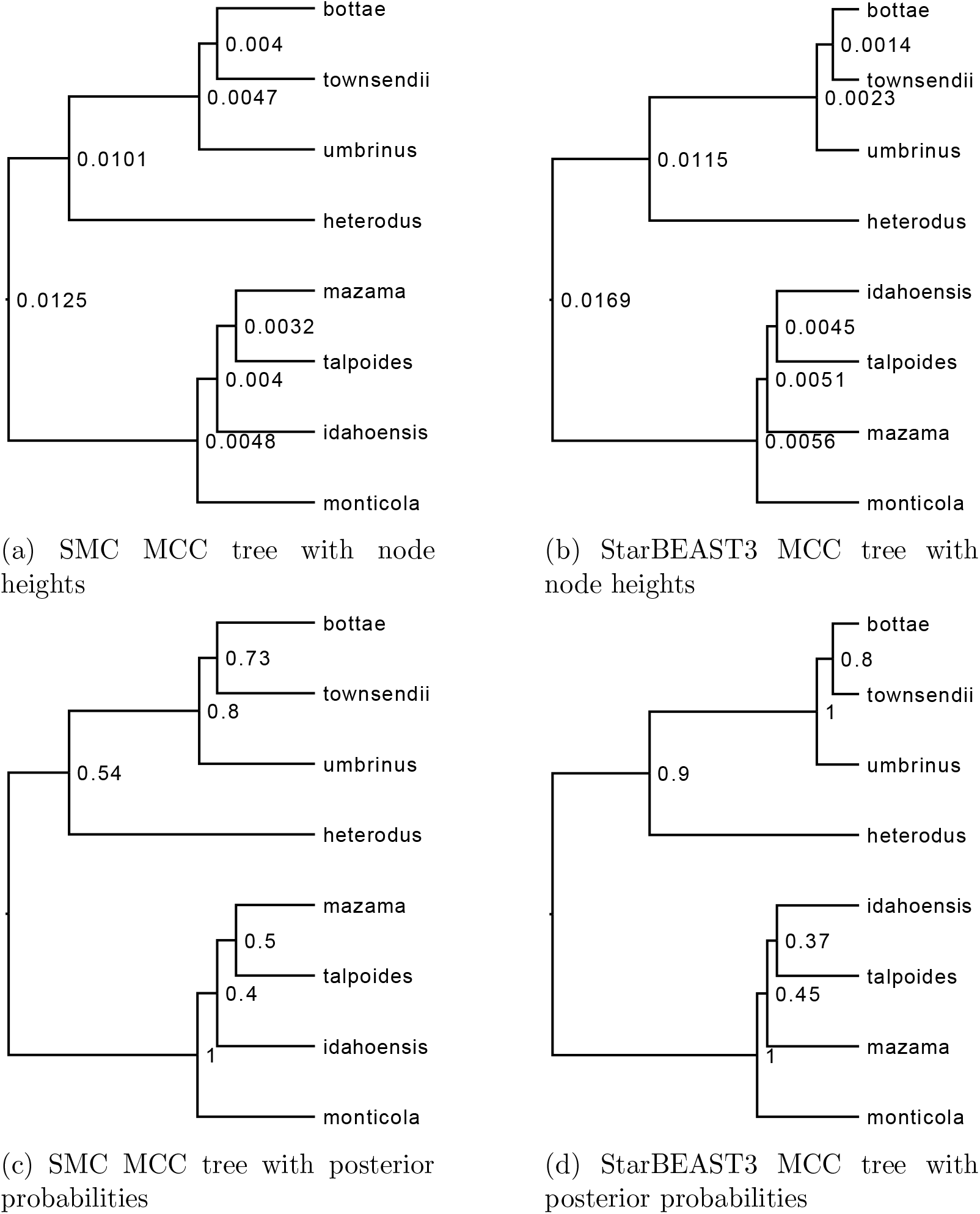
Gopher dataset. a) Maximum clade credibility tree estimated by SMC with node heights. b) Maximum clade credibility tree estimated by StarBEAST3 with node heights. c) Maximum clade credibility tree estimated by SMC with posterior probabilities. d) Maximum clade credibility tree estimated by StarBEAST3 with posterior probabilities. Figures created with TreeAnnotator (Drummond et al., 2007).

### Snake dataset

All methods recovered the same topology, with high support values for the two major clades: *S. edwardsii, S. tergeminus*, and *S. catenatus*; and *S. miliarius, S. barbouri*, and *S. streckeri*. Without accounting for parallelization, SMC took 27411 seconds on average, and StarBEAST3 took 40603 seconds on average.

Maximum clade credibility (MCC) trees (Fig. 9a,b) for SMC and StarBEAST3 show comparable branch lengths between programs. SMC found slightly longer branch lengths for the clade containing *S. barbouri, S. miliarius*, and *S. streckeri*, but branch lengths on the MCC trees for this clade are no more than 0.0039 units different.

**Figure 8:**
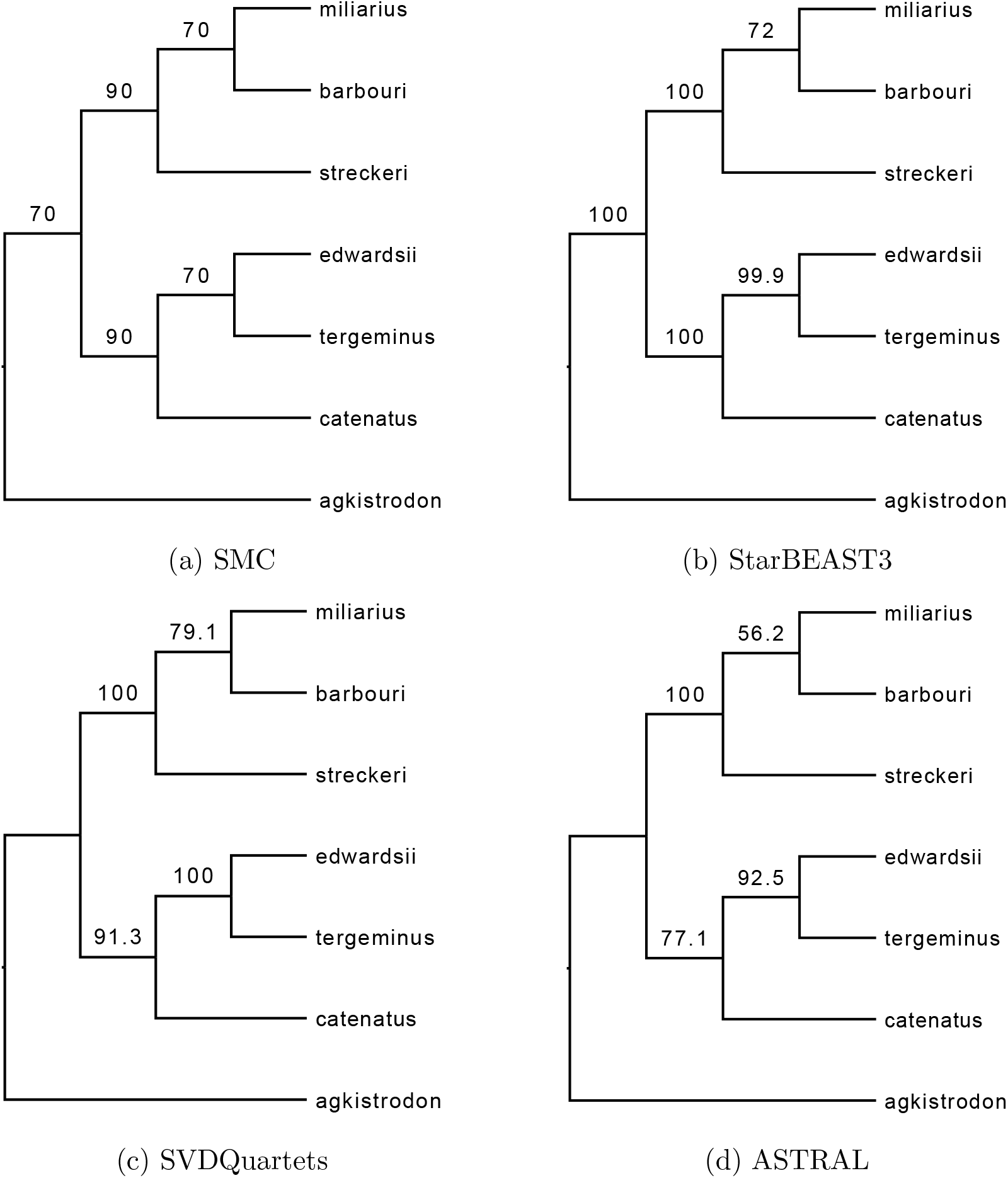
Snake dataset. a) 50% majority rule consensus tree estimated by SMC. b) 50% majority rule consensus tree estimated by StarBEAST3. c) Bootstrapped consensus tree estimated by SVDQuartets. d) Tree estimated by ASTRAL with local posterior probabilities. SVDQuartets and ASTRAL trees rooted at outgroup *Agkistrodon spp*.

**Figure 9:**
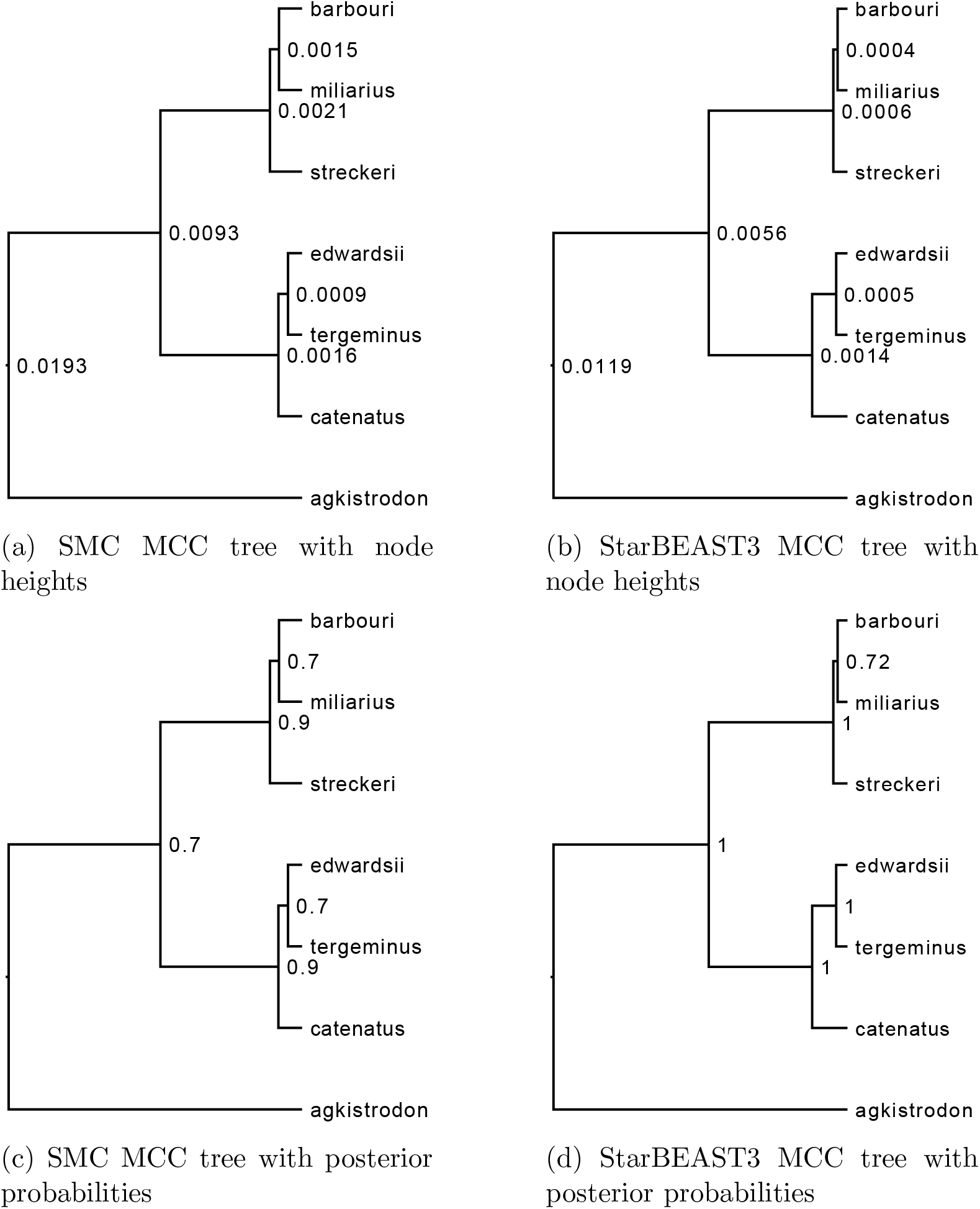
Snake dataset. a) Maximum clade credibility tree estimated by SMC with node heights. b) Maximum clade credibility tree estimated by StarBEAST3 with node heights. c) Maximum clade credibility tree estimated by SMC with posterior probabilities. d) Maximum clade credibility tree estimated by StarBEAST3 with posterior probabilities. Figures created with TreeAnnotator (Drummond et al., 2007).

Posterior probabilities for the MCC trees (Fig. 9c,d) are also comparable. SMC found slightly lower support for *Agkistrodon spp*. as sister to the ingroup taxa and slightly lower support for the group *S. edwardsii* and *S. tergeminus*, but the topologies of both programs are identical.

## Discussion

It is becoming increasingly important for phylogenetic methods to be able to accommodate datasets with hundreds of nuclear loci that conflict with one another due to incomplete lineage sorting or other factors. We have described a fully-Bayesian MSC method that is parallelizable in ways that are not possible for existing multispecies coalescent MCMC algorithms. Bayesian methods sample joint posterior distributions of continuous parameters (e.g., edge lengths, population sizes) and marginalize over discrete tree topologies, which makes them useful even as faster, non-Bayesian alternatives continue to be developed (Chifman and Kubatko, 2014; Mirarab and Warnow, 2015). Bayesian methods are also flexible with respect to model and can be used, for example, to estimate divergence times (Ogilvie et al., 2022), though our SMC implementation does not currently do this.

Speed and accuracy are important metrics for determining the utility of a program, and we have found that, while SMC cannot compete with ASTRAL and SVDQuartets in terms of speed, its accuracy and computational efficiency are comparable to that of Star-BEAST3, currently recognized as the state of the art, even when parallelization is not taken into account. It is difficult to compare speed with parallelization between programs. While StarBEAST3 can be parallelized by placing loci on different processors, there is no speed advantage to using more processors than the number of loci. In contrast, SMC can parallelize across loci and particles, enabling it to take advantage of any number of processors. It is difficult to compare the computational efficiency of programs that are so divergent in their approach. Probably the best unit to use is the number of partial likelihood array calculations; however, we chose the simpler route of choosing settings for both programs that yielded runs of approximately the same total user seconds, giving StarBEAST3 slightly more time than SMC.

Particle degeneracy is a common problem with SMC-based algorithms (Truszkowski et al., 2023), and our SMC approach is no exception. If all samples from a locus have the same topology at the end of the first level SMC, the second level may produce inflated support values for clades in the species tree. We found that doing multiple independent SMC replicates and combining samples produced results comparable to StarBEAST3 for both empirical datasets. While this does increase the computational budget, multiple replicates can be run in parallel, and multiple independent StarBEAST3 runs are also recommended for assessing convergence. We found that particle degeneracy was less of an issue in our simulations, likely because the datasets were smaller and simulated from the exact model used for analysis.

Assessing convergence and the number of particles required in SMC algorithms is difficult. Bouchard-Côté (2012) suggested using effective sample size (ESS) to determine the number of particles needed; if ESS is low, it may mean more particles are needed. However, we have found that there are nearly always SMC steps for which the ESS is very low, regardless of how many particles are used. If the likelihood of one particular join in a gene tree is much greater than that of any other possible join, then any particle that gets this join correct will enjoy a selective sweep. This is as it should be; the low ESS in this case is a consequence of the high marginal gene tree clade posterior. Correctly diagnosing the causes of low ESS remains an area for future work.

We have found that increasing the number of loci in a dataset increases the number of SMC steps but does not necessarily increase the number of particles needed in the first level because only one locus is addressed during each step. Increasing the number of individuals sampled increases the number of particles required in the first-level SMC. This is because more individuals means more join possibilities, especially at early steps or when deep coalescence is common. For example, sampling 100 individuals for a particular species requires 14828 particles to have a 95% chance of getting the first join correct (assuming that the first coalescence event is shallow). Finally, increasing the number of species also increases the number of particles needed because of the smaller chance that any one particle proposes a correct species tree join. This is not surprising because MCMC approaches also require more effort for larger sample sizes (loci, species, individuals per species).

Sampling the multispecies coalescent using SMC thus retains the advantages of fully Bayesian methods and is parallelizable in ways that Bayesian MCMC methods are not but also adds unique challenges. We demonstrated the performance of SMC compared to other commonly-used species tree methods using two empirical datasets and 400 simulated datasets.

## Acknowledgements

We would like to thank Laura Kubatko, Elizabeth Jockusch, Kent Holsinger, and Jill Wegrzyn for their constructive comments.

